# Experimental Estimate of Soil Nutrient Exchange in an Afrotropical Forest: The Role of Dung Beetle Community Complexity

**DOI:** 10.1101/2020.09.29.319327

**Authors:** Roisin Stanbrook

## Abstract

Despite recognition of its importance, little is known about functional aspects of soil fauna. Here, we investigate the effect that different dung beetle functional groups have on macronutrient movement (N, P, K, and C) from dung into soil over 112-day period. We report a large overall effect where more macronutrients are moved into soil over time when beetles are present compared to a control treatment. We also report a large effect of beetle functional groupings on the amount of macronutrient movement, with larger dung beetles moving more nutrients over time. We provide the first experimental evidence that dung beetle body size directly influences macronutrient recycling and discuss the importance of dung beetle functional characteristics in maintaining soil fertility.

## Introduction

There is universal recognition that soil nutrient recycling is fundamental to the maintenance of global ecosystem services. It has been suggested that soil be viewed as natural capital that contributes to the function of ecosystems by maintaining the bioavailability of nutrients and physical structure of the environment (de Groot, Wilson, & Boumans, 2002; Dominati, Patterson, & Mackay, 2010), as well as contributing to human and ecological food security (Barrios, 2007). There is much evidence that soil contributes to the maintenance of biodiversity and stability, for example through the regulation of the microclimate and the control of pathogens (Altieri, 1999). However, while there is wide acknowledgement of the importance of soil systems in contributing to these environmental functions, soil arthropods have received relatively little research attention in comparison to microbes (Lavelle et al., 2006). Further, although the function and importance of dung in nutrient cycling is almost unstudied it is likely to have a critical role in soil environments. This is because most herbivores use only a small proportion of the nutrients they ingest; in mammals, 60-99% of the ingested nutrients returned to the soil in the form of dung and urine (Williams & Haynes, 1990).

One important group in soil nutrient cycling is the paracoprid (tunneling) dung beetles (Nichols et al., 2008). Paracoprids dig tunnels below dung and they bury brood balls in nests which they construct. Incidental nutrient cycling occurs when dung is mechanically relocated underground during nest building. This manipulation is thought to accelerate nutrient breakdown and incorporation of macronutrients, such as faecal nitrogen, directly into the soil (Gillard, 1967; Kakkar, Singh, & Mittal, 2008). This recycling of nutrients has been shown, experimentally, to increase pasture productivity through the incorporation of organic matter into substrates (Bang et al., 2005; Yoshihara & Sato, 2015).

Ecologically, dung beetles have been classified into functional guilds based on traits such as body size, reproductive strategy, flight activity patterns and dung removal behaviour (Doube, 1990; Pincebourde, 2005). There is some evidence that large dung beetles remove greater quantities of dung from soil surfaces (Bui, Ziegler, & Bonkowski, 2020; Frank, Hülsmann, Assmann, Schmitt, & Blüthgen, 2017; Nervo, Tocco, Caprio, Palestrini, & Rolando, 2014), the functional relationship between paracoprid dung beetle trait diversity (e.g. body size or nesting behaviour) and maintenance of soil nutrients due to nutrient recycling remain unclear.

We investigated the effect of functional variation in paracoprid dung beetles on soil macronutrient recycling in an equatorial African ecosystem. We focussed on the ecological effect on nutrient cycling in paracoprid tunnellers, as they comprise the most abundant functional dung beetle group (Adrian L.V Davis, Frolov, & Scholtz, 2008) and have previously been shown to have a large role in dung removal (Slade et al., 2007). Our overall aim was to test whether there is a strong functional effect of dung beetle body size on the quantity of macronutrients passed from elephant (*Loxodonta africana*) dung into the soil. Specifically, our objectives were to 1) assess whether the transfer of nutrients from dung to soils is influenced by dung beetle body size, and 2) estimate the temporal effect of the dung beetles on dung to soil nutrient transfer. We discuss our findings in the context of the functional diversity of soil macrobiota and its implications for soil nutrient enrichment.

## MATERIALS AND METHODS

The study was conducted within the Aberdare National Park (ANP), Nyeri County, Kenya (0.4167° S, 36.9500° E). The ANP is ring-fenced and is contained within the forested Aberdare range, which is an elongated mountain range, running approximately north south, parallel to the direction of the Rift Valley, 60 km to the west of Mt. Kenya. The slopes are steep and densely forested while the foothills have been cleared of forest and are intensively farmed with crops such as pyrethrum and coffee (Lambrechts, Woodley, Church, & Gachanja, 2003). The underlying soil is mollic Andosol (Mugendi, Mucheru-Muna, Mugwe, Kung’u, & Bationo, 2007), a part-volcanic, humus rich, gritty loam with a high level of phosphorus absorption but low levels of phosphate availability (Orgiazzi et al., 2016). This means that although the retention of phosphorus is relatively high in comparison with other soil types due to the low cation exchange capacity very little of this is available for plant uptake.

### Functional Guilds

Dung beetles were collected using dung baited pitfall traps 24 hours before the start of the experiment. All captured individuals were identified to genus. Total body length (anterior clypeal sinuation to pygidium) was measured to the nearest millimetre using digital callipers, and mass was measured to the nearest gram. Functional guilds can be defined as groups of species which possess a suite of traits allowing them to exploit the same resource in a similar way (Stroud et al., 2015). Dung beetles were classified into functional guilds using functional guild categories identified in Doube (1990) that are based on body size, then assigned to one of three treatments (see Table 1): (1) small (body size range: >5mm to <15mm), (2) medium (>15mm to <25mm), or (3) large (>25mm). We also had a negative control treatment with no beetles. Each treatment type contained an equal biomass of beetles (8.1 ± 0.04 grams).

| Treatment | Body Size (mm) | Genera | Total Number of Individuals *per* Genera | Proportion in Treatment (%) |
| --- | --- | --- | --- | --- |
| Large | >25mm | *Heliocopris* | 2 | 100 |
| Medium | >15 mm <25 | *Onitis* | 8 | 80 |
|  |  | *Diastellopalpus* | 4 | 15 |
|  |  | *Copris* | 6 | 5 |
| Small | >5 mm <15mm | *Onthophagus* | 16 | 70 |
|  |  | *Milichus* | 22 | 10 |
|  |  | *Oniticellus* | 6 | 12 |
|  |  | *Liatongus* | 8 | 6 |
|  |  | *Euoniticellus* | 12 | 1 |
|  |  | *Caccobius* | 6 | 1 |

### Mesocosm design

Two replicates of four treatment types were used to assess macronutrient transfer. Each treatment was represented by an experimental mesocosm containing exclosures which contained dung and the dung beetles. Each exclosure consisted of a 40L (height: 50cm × 40cm) plastic bucket buried with the top lip placed flush with the soil surface. 40L of excavated soil was sifted with a 2mm aperture to remove debris and macroinvertebrates and was then placed back into each bucket until it was completely filled. Freshly deposited elephant dung was collected from the top section of boli, leaving behind dung that was in contact with the ground to avoid soil contamination. Similarly, dung contaminated by urine was not used. Dung was shaped into hemispherical 1L pats and frozen for 20 hours to kill any macroinvertebrates present. Dung pats were defrosted at room temperature and placed on top of each soil-filled bucket and then the dung beetles for each treatment type were released. A pyramidal structure made of wooden poles wrapped in 1.2mm gauge netting was placed above each bucket to prevent ingress or egress of dung beetles during the experiment. In the control treatment, a dung pat was placed but no beetles were released. The experiment began on the 28 April 2015 and ended after 112 days, as this timeframe covered the expected completed lifecycle for all species used in the experiment and allowed the action of both adult and larval dung beetles to be recorded.

### Soil samples

Soil samples were collected by using a standard soil corer (2.5 × 10cm) with one core collected from under each pat at the start of the experiment (day 0) and subsequently at days 7, 14, 28, 56 and 112. Each soil sample was frozen at −20°C in preparation for transport and laboratory analysis. Dung samples were dried for 24hrs at 70°C, then pulverized in a ceramic mortar to pass through a 2mm sieve and were analysed for Nitrogen (N), Phosphorus (P), and Potassium (K) concentrations using the Mehlich-3 extraction procedure (Mehlich, 1984). We added five grams of dung to 20ml of 0.05 M HCl in 0.025 M H2SO4, and the filtrate was analysed by Inductively Coupled Plasma-atomic Emission Spectrometry (ICP-OES). Approximately 5g of dried and weighed soil were decarbonised with 1M solution of HCl before being analysed for total C and N concentrations through the LECO TruSpec analyser using the combustion (Dumas) method. Data were analysed using a linear mixed effects approach with time and Functional Group as explanatory variables and nutrient transfer as the response variable. All analysis were completed using the nlme package (Pinheiro et al., 2017) in R software version 3.1.1. (R Core Team, 2016).

## Results

There was a highly significant effect between treatments for all tested macronutrients across the 112-day experimental period (all P < 0.05 for C, N, P and K, see Table 2; Figure 1). That is, the presence of beetles in our treatments increased nutrient uptake in the soil for all treatments, relative to passive leeching of nutrients from dung in the absence of beetles in our control treatment. Large-bodied beetles effected the greatest change in macronutrient status, enriching the soil on average by 44.51% for all macronutrients in comparison to the control treatments. All functional groups had a significant effect on available P transfer from dung into the soil. The available P content in each treatment increased rapidly from day 0 for all functional guilds but appeared to stabilise by day 56 of the study (Figure 1C).

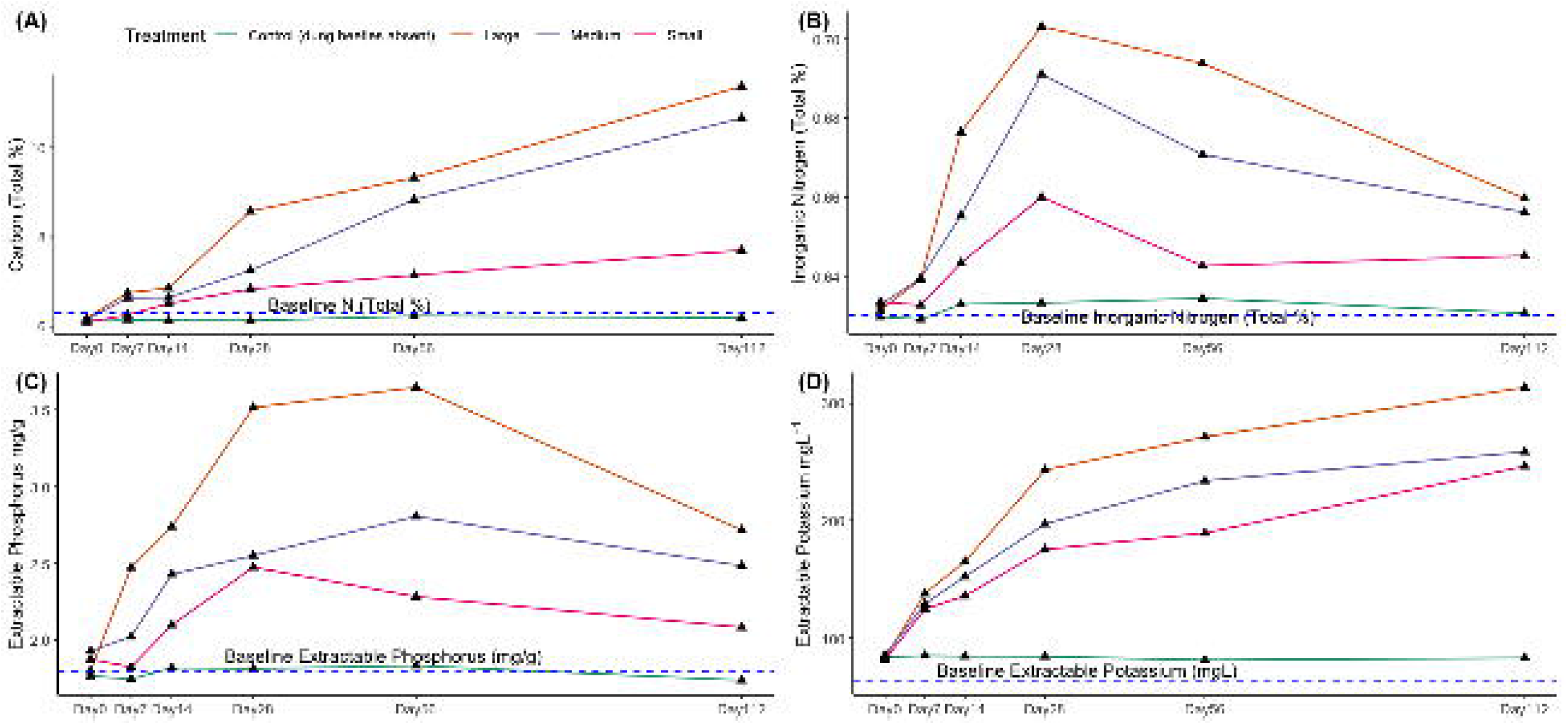

| Treatment main effect |  |  | Treatment effect over time |  | Large |  | Medium |  | Small |  |
| --- | --- | --- | --- | --- | --- | --- | --- | --- | --- | --- |
|  | F (df) | *P* | F (df) | *P* | contrast | *P* | contrast | *P* | contrast | *P* |
| N | 4.61 (3, 15) | **0.01** | 8.31 (3, 39) | **0.001** | 0.26 | **0.002** | 0.13 | 0.08 | 0.09 | 0.1 |
| C | 4.84 (3, 15) | **0.01** | 10.16(3, 39) | **0.01** | 3.14 | **0.002** | 2.56 | **0.01** | 1.67 | 0.0 |
| P | 10.68 (3, 15) | **<0.001** | 0.36 (3, 39) | **<0.05** | 1.63 | **<0.001** | 1.21 | **0.003** | 0.75 | **0.0** |
| K | 14.17 (3, 15) | **<0.001** | 21.76 (3, 39) | **<0.001** | 118.72 | **0.001** | 92.54 | **<0.001** | 75.33 | **0.0** |

When between treatment effects were analysed, the greatest effects were observed between the control (dung + no beetles) and the large body size functional guild (beetles with a body length >0.25mm; Table 2). Thus, large bodied beetles accounted for the greatest transfer of nutrients into the soil for all macronutrients we measured; Carbon (P=0.002); Nitrogen (P=0.002); Potassium (P=0.0001); Phosphorus (P<0.001) over time with the largest overall effect for the transfer for exchangeable Potassium (Figure 1). The small-bodied functional guild showed the smallest effect for macronutrient transfer to the control; with significant effects for K (P=0.003) and P (P=0.01), but not for N or C (both P>0.05; see Figure 1 and Table 2). The medium-bodied functional guild showed a moderate effect on soil macronutrient enrichment with significant effects for K (P=0.0008), P (P=0.003), and C (p=0.01), but no difference for N (P=0.08).

## Discussion

Our main finding was that paracoprid dung beetles have a large and positive effect on the incorporation of macronutrients from dung into the soil and that this effect increases over time to a period of at least 112 days (Figure 1). When the functional guilds we identified are ranked in order of their capacity to facilitate nutrient exchange, large beetles have the largest effect, followed by medium and small-bodied beetle groups, respectively. Our results also suggest the movement process and rate from dung to soil differed *per* nutrient. Inorganic N content in the soil from all the treatments increased from day 14 for all treatments and tended to increase again from days until 56 where they tapered off. Inorganic N content in the soil only significantly increased in the presence of the large-bodied functional guild. The transfer of readily available K content was much faster than those of other nutrients, irrespective of the dung beetle treatments. Hogg, (1981) reported that most K in dung is water soluble and that the contents of water-soluble N and P in dung are relatively small. Therefore, the difference in movement of those nutrients from the dung to the soil are possibly explained by the difference in their water solubility.

The largest beetles in our experiment are in the genus *Heliocopris*, which contains species that are among the largest dung beetles in the world (Pokorný, Zidek, & Werner, 2009) and are known for their ability to relocate large quantities dung underground (Kingston & Coe, 1977; Klemperer & Boulton, 1976; Stanbrook, in press). They tend to specialize on the dung of large herbivores such as elephant and rhino. *Heliocopris* occurs at relatively low population density, most likely because of their large body size and the low density of their preferred dung type (Davis, 2013). However, their large body size is frequently cited as a trait that correlates significantly with increased extinction risk over ecological time scales and has been reported in both vertebrates and invertebrates. Indeed, several studies highlight declines in large-bodied dung beetles in the presence of habitat disturbance (Gardner, Hernández, Barlow, & Peres, 2007; Larsen, Williams, & Kremen, 2005), and concomitant decline of large herbivore density (Bogoni et al., 2016).

Slade et al. (2007) assessed dung beetle morphological traits in the context of ecosystem functioning and reported that the absence of large, nocturnal tunnellers yielded a 75% reduction in the quantity of dung removed from soil surfaces. Other studies investigating different aspects of functional diversity have established that single species may be more influential in terms of ecosystem services provision than overall species richness. (e.g. Larsen et al., 2005; Soliveres et al., 2016). These observations are congruent with our findings which suggest that the largest dung beetles are, functionally, the most important species in effecting soil nutrient transfer from dung they are more effective at burying larger quantities of dung. However, these large dung beetles, in general, appear to be the least tolerant to habitat perturbation and other drivers of ecosystem change (Séguin et al., 2014) and (Díaz et al., 2013).

Soil nutrient depletion has been linked with declines in crop productivity in sub Saharan Africa, (Sanchez et al., 1997) and Kenya is particularly affected by falling agricultural productivity and diminishing food security, with 12 million people residing in areas with land degradation (Mulinge et al., 2016). Food webs may be linked across habitat boundaries and the biodiversity of one ecosystem, in this case a Protected Area, may influence the functional delivery of services to adjacent ecosystems such as the agriculturally important land described here. Historically, soil fertility depletion is the major biophysical cause of declining crop productivity and a fundamental root cause for declining food security on smallholder farms in central Kenya (Mugendi et al., 2007; Njeru et al., 2011). This may have an impact on a local scale and may indirectly affect the agriculturally dependent communities which surround the Aberdare National Park. This study reinforces the importance of understanding dung beetle functional characteristics and highlights that loss of dung beetle species could negatively impact the transfer of important macronutrients to the soil. It is important therefore to safeguard those species that are the most important for sustaining ecosystem function and ascertain how sensitive they may be to anthropogenic activity.

